# The Treatment of Clear Cell Ovarian Cancer with the Poly(ADP- Ribose) Polymerase (PARP1) Inhibitors (AG14361,Veliparib,Olaparib) as Chemosensitizers

**DOI:** 10.1101/2020.11.10.377424

**Authors:** Joseph Angel de Soto

## Abstract

**Introduction:** Most of those who get ovarian cancer will die from this cancer. Of the major types of ovarian cancer clear cell carcinoma is the most aggressive and chemoresistant type of epithelial ovarian cancer. Here the sensitivity of clear cell ovarian carcinoma to poly adenosine diphosphate [ADP-ribose] polymerase (PARP) inhibitors is tested.

**Methodology:** Ovarian cancer cell lines were treated with the PARP inhibitors AG14361, Veliparib, or Olaparib alone or in combination with cisplatin, carboplatinum, doxorubicin, 5-fluorouracil (5-FU), gemcitabine and paclitaxel for 72 hours. The IC50 concentrations were calculated. Each experiment was replicated 10 times.

**Results:** As single agents the PARP inhibition of ovarian cancer among serous, endometroid and clear cell ovarian cancer cell lines was similar. Clear cell ovarian cancer seemed particularly susceptible to chemo-sensitization by PARP inhibitors with paclitaxel, 5-FU, carboplatin, doxorubicin and/or cisplatin. Antagonism was seen with gemcitabine.

**Conclusion:** PARP inhibitors are exceptional chemosensitizers of clear cell ovarian cancer to treatment with most standard chemotherapy agents.

## Introduction

The American Cancer Society and National Cancer Institute estimates that there will be over 22,240 new cases of ovarian cancer with over 14,000 deaths from ovarian cancer this year.^1^ The median age for diagnosis for sporadic disease is 60 years with those with a genetic tendency towards ovarian cancer often being diagnosed in their 50’s. Ovarian cancer represents tumors of epithelial, germ cell and sex cord-stromal origin. Epithelial ovarian cancer which arises from arise from the surface epithelium of the ovary represents about 90% of ovarian cancers. The most important risk factors for the development of ovarian cancer are age, family history and infertility while lactation, pregnancy, and tubal ligation are mildly protective.^2^ The ovarian epithelium cancers can be divided into four types: papillary serous, endometroid, mucinous and clear cell. Clear cell represents represent 6% of the epithelial cells types and is the most aggressive and resistant to therapy.^3^ Following cytoreductive surgery for ovarian cancer the use of platinum – taxane based chemotherapy has traditionally been the gold standard for chemotherapy.^1^ Although most ovarian cancers are chemosensitive at diagnosis drug acquired resistance frequently appears leading to progression and ultimately death.

The hallmark of cancer development is the development of clonal expansions genetic mutations which continue to accrue additional genetic mutations and eventually genomic instability. This led to the idea that perhaps one could take advantage of this genetic instability in cancer cells an instability not present in non-cancer cells.^4^ Thus, providing a pathway to targeted therapy. In a paper by Farmer that appeared in Nature, the feasibility of using PARP inhibitors as targeted therapy was shown.^5^ It has reported that cells with BRCA1 or BRCA2 mutations which are important for homologous recombination were 1000 times more sensitive to PARP inhibition than those cell without a defect in homologous recombination.^6^ Papers by De Soto and another by Calabrese illustrated the clinical feasibility of using PARP inhibitors as chemosensitizers in the treatment of human cancer.^7,8^ Poly (ADP-Ribose) Polymerase (PARP) is a key enzyme utilized in the repair of single strand DNA damage via the Base Excision Repair pathway. PARP inhibition produces persistent single-strand DNA breaks which during mitosis become double stranded breaks at the replication fork.^9^ These double stranded breaks may be repaired by homologous recombination. However, many cancers are deficient in their ability to repair by homologous recombination due to deficiencies in BRCA1, BRCA2 and other enzymes needed in this repair pathway. This leads to the accumulation of chromosomal damage and the death of cancer cells. Because most non-cancer cells are efficient in homologous recombination, they are relatively unaffected by the inhibition of PARP. Myelosuppression however dies occur with PARP inhibitors in addition to somnolence.^10,11^ Today, PARP inhibitors are approved for maintenance therapy in ovarian cancer following platinum based therapy.^12^ PARP inhibitors are also treatment friendly as Olaparib and Veliparib both have the advantage of being available orally.^13, 14^ Here we show the effective use of PARP inhibiters as chemosensitizers against clear cell carcinoma with taxane, platinum and other agents used to treat ovarian cancer.

## Materials & Methods

### Cell Lines & Medications

ES-2, TOV-112D, TOV-21G, and SKOV-3 and human epithelial ovarian cancer cell lines where obtained from American Type Tissue Culture in Manassas VA. Cisplatin, Carboplatinum, Doxorubicin, 5-Fluorouracil, Gemcitabine and Paclitaxel where purchased from Sigma-Aldrich, St. Louis MO. The PARP-1 inhibitor AG14361 was obtained from Organix inc. in Woburn MA while ABT888 (Veliparib) and AZD2281 (Olaparib) were obtained from Selleck Chemical in Houston TX. Cells were grown in DMEM media with 10% fetal bovine serum in a 5% CO_2_ incubator at 37 °C. Cells at 80-85% confluence were trypsinized, washed with PBS and plated for each experiment

### Determining IC50 Values

Standard curves where made by plating 25k to 300k cells for each cell line for 4-12 hours and after ascertaining that the cells where attached exposed to a solution of 1 mg/ml thiazolyl blue tetrazolium (MTT) for 30 minutes. This was followed by decanting the MTT and adding propranolol to each well for 30 minutes. The absorption for each well was read with the Perkin-Elmer 1420 multi-label counter. Each data point was replicated at least 10 times. IC50 values were then determined by plating cells in 12 well plates and after incubation for 4-12 hours the cells were exposed for 72 hours to various doses of drug. There were at least two controls containing vehicle for each experiment. The number of cells in each well at each specific drug concentration was then determined by the previously described MTT assay. Each experiment was replicated at least 9 times with the tenth time being a direct cell count by hemacytometer. The IC 50 values were then calculated by graphing the % inhibition vs log(dose) using Sigma Plot and standard 3rd order (sigmoidal) equations with the required r2 value having to be > 0.975.

The “PARP in vivo Pharmacodynamic Assay II kit” from Trevigen inc, Gaithersburg MD was used to determine the amount of PARP activity in each cell line prior to and after treatment with PARP inhibitors.

## Results

The IC50 values of AG14361, Veliparib and Olaparib against clear cell, serous and mucinous epithelial ovarian cancer cell lines was evaluated. The ES-2 clear cell ovarian carcinoma was derived from a 47 year old African American female with low to moderate level resistance to cisplatin and doxorubicin.^15^ This clear ovarian cancer was very sensitive to PARP inhibition with the IC50 values for AG14361, Veliparib and Olaparib being 6.2 μM, 38 μM and 97 μM. A second clear cell ovarian carcinoma TOV21G was derived from a 62 year old female was also found to be very sensitive to PARP inhibition.^16^ The following are the IC50 values of the PARP inhibitors against the TOV21G ovarian cancer cell line: AG14361 8.9 μM, Veliparib 78 μM and Olaparib 150 μM. A serous ovarian cancer cell line SKOV-3 derived from a 64 year old female with tumor resistance to cisplatin and doxorubicin was tested against AG14361, Veliparib and Olaparib with the IC50 values being 15.2 μM, 82 μM and 17.6 μM. The Ov-90 cell line was another serous cell ovarian cancer cell line evaluated which was derived from a 64 year old French female.^17^ The OV-90 cell line is Her2/neu + and like the clear cell line TOV-21G has a deletion at 3p24. Interestingly, this same locus has been found to be associated with breast cancer. The IC50 values for the Ov-90 cell line were 5.2 μM, 35 μM and 15 μM for AG14361, Veliparib and Olaparib respectively. Next the ability of the PARP inhibitors to inhibit endometrioid type ovarian cancer was evaluated through the TOV112D ovarian cancer cell line. This cell line was derived from a 42 year old French female and was her2/neu +.^16^ The IC50 values after exposure to the PARP inhibitors were similar to those obtained for the serous cell lines the results are summarized in Fig I.

Evidence suggests that the PARP inhibitors may be able to chemo-sensitize breast and brain cancers to DNA damaging agents. The IC50 values for five commonly used chemotherapeutic agents used to treat ovarian cancer cisplatin, carboplatin, doxorubicin, 5-fluorouracil, gemcitabine and paclitaxel where determined for the ES-2, TOV112D and TOV21G cell lines (Table 2). Next, we determined the IC50 value of these same agents when used in combination with a constant 10 μM of AG14361. The value of 10 μM was chosen as it is a blood level which represents a blood level which might clinically be reached not only for AG14361 but Veliparib and Olaparib, thus potentially allowing for the translation of these results into the clinic.

**Table I.** The IC50 values of selected PARP inhibitors against sporadic epithelial ovarian cancer.

| **Cell Line**<br>**Subtype** | **AG14361**<br>**IC<sub>50</sub>** | **Veliparib**<br>**IC<sub>50</sub>** | **Olaparib**<br>**IC<sub>50</sub>** |
| --- | --- | --- | --- |
| **TOV21G**<br>**Clear Cell** | **8.9 µM** | **78 µM** | **150 µM** |
| **ES-2**<br>**Clear Cell** | **6.2 µM** | **38 µM** | **97 µM** |
| **SKOV-3**<br>**Serosus** | **15.2 µM** | **82.0 µM** | **17.6 µM** |
| **OV-90**<br>**Serosus** | **5.2 µM** | **35.0 µM** | **15.0 µM** |
| **TOV112D** | **14.6 µM** | **63.8 µM** | **32.4 µM** |
| Endometrioid |  |  |  |

**Table 2.** The IC50 values of selected chemotherapy agents against sporadic epithelial ovarian cancer.

|  | ES-2<br>Clear cell | TOV112D<br>Endometrioid | TOV-21G<br>Serous |
| --- | --- | --- | --- |
| Cisplatin | 1.18 μM | 5.4 μM | 17.5 μM |
| Doxorubicin | 0.046 μM | 0.038 μM | 0.082 μM |
| 5-Fluorouracil | 20.7 μM | 19.9 μM | 40. 5 μM |
| Gemcitabine | 0.00012 μM | 0.0046 μM | 8.6 μM |
| Paclitaxel | 0.018 μM | 0.008 μM | 0.075 μM |

**Table 3.**

| PARP inhibitor + Chemotherapeutic Agent |  |  |  |  |  |  |
| --- | --- | --- | --- | --- | --- | --- |
| ES-2 Ovarian Cancer Cell Line (IC50 values) |  |  |  |  |  |  |
| Drug | + 10 μM<br>AG14361 | $\Delta$ | + 10 μM<br>Veliparib | $\Delta$ | +10 μM<br>Olaparib | $\Delta$ |
| Cisplatin | 0.002 μM | 590x | 0.01 μM | 118x | 0.005 μM | 235x |
| Carboplatin | 0.001 μM | 18,800x | 11.2 μM | 1.7x | 1 μM | 18.8x |
| Doxorubicin | 0.00023 μM | 200x | 0.017 μM | 2.7x | 0.009 μM | 5.1x |
| 5-Fluorouracil | 0.001 μM | 20,700x | 6.1 μM | 3.4x | 20.2 μM | 1.02x |
| Gemcitabine | 0.00012 μM | 1x | 0.0028 μM | 0.04x | 0.0022 μM | 0.054x |
| Paclitaxel | 0.018 μM | 180x | 0.043 μM | 1.4x | 0.0001 μM | 600x |
| TOV-112D Cancer Cell Line (IC50 values) |  |  |  |  |  |  |
| Cisplatin | 0.1 μM | 54x | 3.8 μM | 1.4x | 1.0 μM | 5.4x |
| Carboplatin | 42.0 μM | 1.7x | 70.0 μM | 1x | 50.0 μM | 1.4x |
| Doxorubicin | 0.013 μM | 2.9x | 0.018 μM | 2.1x | 0.031 μM | 1.2x |
| **5-Fluorouracil** | **1.0 μM** | **19.9x** | **5.9 μM** | **3.4x** | **8.0 μM** | **2.5x** |
| **Gemcitabine** | **0.03 μM** | **0.11x** | **0.01 μM** | **0.33x** | **0.038 μM** | **0.86x** |
| **Paclitaxel** | **0.0008 μM** | **25x** | **0.013 μM** | **1.4x** | **0.0012 μM** | **15x** |
| **TOV21G (IC50 values)** |  |  |  |  |  |  |
| **5-Fluorouracil** | **0.001 μM** | **40,500x** | **6.8 μM** | **6x** | **28.0 μM** | **1.4x** |
| **Paclitaxel** | **0.005 μM** | **14x** | **0.005 μM** | **4.4x** | **0.014 μM** | **1.5x** |

## Discussion

It has been shown that up to 50% of ovarian cancers may have trouble repairing double strand breaks via homologous recombination.^18^ The addition of adjuvant chemotherapy in clear cell carcinoma does not show any survival difference following surgical excision in stage I and stage II cancer and little improvement in stage III or IV.^19^ Thus, the need for improved cytotoxicity of adjuvant chemotherapy towards clear cell ovarian cancer is needed. Clear cell ovarian cancer has been compared to triple negative breast cancer due to its absence of both estrogen and progesterone receptors and rapid growth.^20^ Dysfunctional homologous recombination is a determinant in platinum and PARP sensitivity and possibly taxane sensitivity providing a rational for the use of PARP inhibitors in conjunction with these other agents in treating ovarian cancer.^21^ Loss of the ability to perform homologous recombination may be due to either genetic mutation or epigenetic influences.^22^ The increased amount of double stranded breaks leads to rapid chromosomal instability and cellular death. Among the proteins that may have mutations or be epigenetically silenced are BRCA1, BRCA2, PALB2, ATM, CHEK1, CHEK2, and RAD51.^23^ The loss of homologous recombination is also associated with TP53 mutation and hence, a loss of cell cycle regulation.^24^ Which may be important in clear cell ovarian cancer which is an exceptionally fast growing cancer. Additionally, PARP may act as a chemosensitizer by also partially disrupting spindle assembly.^25^ PARP inhibitors may be especially useful for those cancers that are resistant to prior platinum and other treatments but, are limited in their ability to chemosensitize these agents in cancers that are refractory to specific chemoagents.^26^ Refractory behavior is often seen in ovarian cancer that has a back mutation in previously dysfunctional enzymes needed for homologous recombination.^27^ The use of PARP inhibitors may also be useful in treating those clear cell carcinomas that are cross resistant to platinum, topoisomerase and ionizing radiation therapy providing a grim outcome for these patients. This type of cross resistance is often due elevated glutathione-S-transferase which detoxifies these agents.^15^ The use of PARP inhibitors would allow for a cytotoxic effect for the reduced levels of chemotherapeutic agents.

The use of PARP inhibitors maybe useful as single agent maintenance therapy in clear cell carcinoma as the sensitivity to PARP is at least as great as serous cell ovarian cancer which has been shown to be amiable to maintenance therapy.^28^

## Conclusion

The use of PARP inhibitors to treat clear cell ovarian cancer in conjunction with platinum agents, taxane or topoisomerase inhibitors may allow for using a lower dose of these toxic agents and thus reduce the side effects. Additionally, the use of PARP inhibitors as chemosensitizers may allow for the more targeted and efficient killing of clear cell ovarian cancer cells. PARP inhibitors are exceptional chemosensitizers of clear cell ovarian cancer to treatment with most standard chemotherapy agents. Gemcitabine however is however antagonized by PARP inhibitors. This is the first paper to specifically show the effectiveness of PARP inhibitors against clear cell ovarian cancer.

## Acknowledgments

Bonnie Grunther for proofreading the manuscript

## Conflicts of Interests

None

## Notes

### Competing Interest Statement

The authors have declared no competing interest.

